# Peri-saccadic perceptual mislocalization is different for upward saccades

**DOI:** 10.1101/333112

**Authors:** Nikola Grujic, Nils Brehm, Cordula Gloge, Weijie Zhuo, Ziad M. Hafed

## Abstract

Saccadic eye movements, which dramatically alter retinal images, are associated with robust peri-movement perceptual alterations. Such alterations, thought to reflect brain mechanisms for maintaining perceptual stability in the face of saccade-induced retinal-image disruptions, are often studied by asking subjects to localize brief stimuli presented around the time of horizontal saccades. However, other saccade directions are not usually explored. Motivated by recently discovered asymmetries in upper and lower visual field representations in the superior colliculus, a structure important for both saccade generation and visual analysis, here we observed significant differences in peri-saccadic perceptual alterations for upward saccades relative to other saccade directions. We also found that, even for purely horizontal saccades, perceptual alterations differ for upper versus lower retinotopic stimulus locations. Our results, coupled with conceptual modeling, suggest that peri-saccadic perceptual alterations might critically depend on neural circuits, like superior colliculus, that asymmetrically represent the upper and lower visual fields.

## Introduction

Saccades are instrumental for rapidly allocating foveal visual processing resources to interesting scene locations. However, each saccade introduces spurious shifts in retinal images, which normally go perceptually unnoticed, and the mechanisms underlying such perceptual stability are an active area of research (Bremmer et al. 2009; Brenner et al. 2008; De Pisapia et al. 2010; Duhamel et al. 1992; Hafed 2013; Hafed et al. 2015; Maij et al. 2010; Maij et al. 2011; Rao et al. 2016; Richard et al. 2009; Ross et al. 2001; Sommer and Wurtz 2008; Watson and Krekelberg 2009; Wurtz 2008; Wurtz et al. 2011; Ziesche and Hamker 2014; Zirnsak and Moore 2014).

Experimentally, perceptual stability mechanisms are studied by peri-saccadically presenting briefly flashed visual stimuli (Ross et al. 2001). If saccades are associated with retinal and/or extra-retinal mechanisms for handling saccade-induced retinal-image disturbances, then perception of such brief flashes may be altered. The flashes effectively allow capturing momentary “snapshots” of the state of the visual system around the time of saccades. In cases with sufficiently low contrast flashes, these flashes may go completely unnoticed in the phenomenon of “saccadic suppression” (Beeler 1967; Burr et al. 1994; Chen and Hafed 2017; Diamond et al. 2000; Hafed and Krauzlis 2010; Zuber and Stark 1966), which has its own rich set of properties and unsolved questions (Bellet et al. 2017; Benedetto and Morrone 2017; Bremmer et al. 2009; Chen and Hafed 2017; Crevecoeur and Kording 2017; Hafed and Krauzlis 2010; Krekelberg 2010; Zanos et al. 2016). However, with sufficiently visible flashes, even though such flashes are easily detected, their locations are grossly misestimated (peri-saccadic perceptual mislocalization) (Cai et al. 1997; Honda 1989; 1991; Lappe et al. 2000; Maij et al. 2010; Morris et al. 2012; Ross et al. 1997; Zirnsak et al. 2014). Under conditions of bright illumination, misestimates of flash location appear as if the flashes are perceptually compressed towards the saccade target (Lappe et al. 2000; Ross et al. 1997), and it is this “perceptual compression” that we sought to explore.

Compression was initially studied along only one dimension and with horizontal saccades (Ross et al. 1997). Subjects made large horizontal eye movements, and flashes could appear along the same direction of the eye movements either at farther or nearer eccentricities than the saccade target. Peri-saccadic flashes were perceptually compressed towards the saccade target such that flashes more eccentric than the target were mislocalized backwards in a direction opposite to the saccade direction and flashes less eccentric were mislocalized forwards. Subsequent studies explored two-dimensional patterns of compression with flashes off the axis of the saccade (Kaiser and Lappe 2004) and discovered asymmetries in two-dimensional compression as a function of flash eccentricity. While the mechanisms behind these asymmetries are not fully understood, existing models agree that they may reflect interactions between oculomotor signals associated with saccade generation, on the one hand, and visual signals associated with flash locations, on the other, and, critically, in sensory and motor neural maps that exhibit “ foveal magnification” (i.e. more neural tissue dedicated to central than peripheral visual field eccentricities) (Hamker et al. 2008; Hamker et al. 2011; Kaiser and Lappe 2004; Richard et al. 2009; VanRullen 2004; Zirnsak et al. 2010).

In this study, we explored this idea further by asking whether peri-saccadic perceptual mislocalization depends on movement direction. Prior experiments, exploring rightward and downward saccades (Kaiser and Lappe 2004; Ross et al. 1997; Zimmermann et al. 2015), did not reveal significant differences in mislocalization patterns as a function of saccade direction. However, our goal was to investigate this question in more detail, and for upward saccades in particular. We were specifically motivated by the notion that if perceptual mislocalization arises as a result of foveal magnification in sensory-motor maps, then other patterns of neural tissue magnification might dictate asymmetries in the patterns of peri-saccadic perceptual mislocalization. For example, if the superior colliculus (SC), as a visual and saccade-related structure (Basso and May 2017; Gandhi and Katnani 2011; Sparks and Mays 1990; Veale et al. 2017; Wurtz 1996), were to be involved in perceptual effects associated with mislocalization (Hafed 2013; Richard et al. 2009; Veale et al. 2017), then this structure’s putative magnification of upper visual field representations (Hafed and Chen 2016) might mean that mislocalization would be different for upward saccades. If so, this would provide important constraints on neural loci and mechanisms for such a highly robust perceptual phenomenon, and would therefore more generally have strong implications on our understanding of perceptual stability in the face of continuous eye movements.

## Materials and Methods

### Subjects

We recruited 4 naïve subjects (2 females; 21-25 years old) in our main experiment using cardinal saccade directions (e.g. see Fig. 1 and **Behavioral tasks** below). We then recruited 2 of the same subjects (1 female) for a second experiment involving oblique saccades (e.g. see Fig. 8 and **Behavioral tasks** below). All subjects provided informed, written consent in accordance with the Declaration of Helsinki, and the experiments were approved by ethics committees at Tübingen University. Each subject completed 5 sessions of 1.5 hours each, with each session consisting of 4 blocks of 160 trials per block.

**Figure 1.**
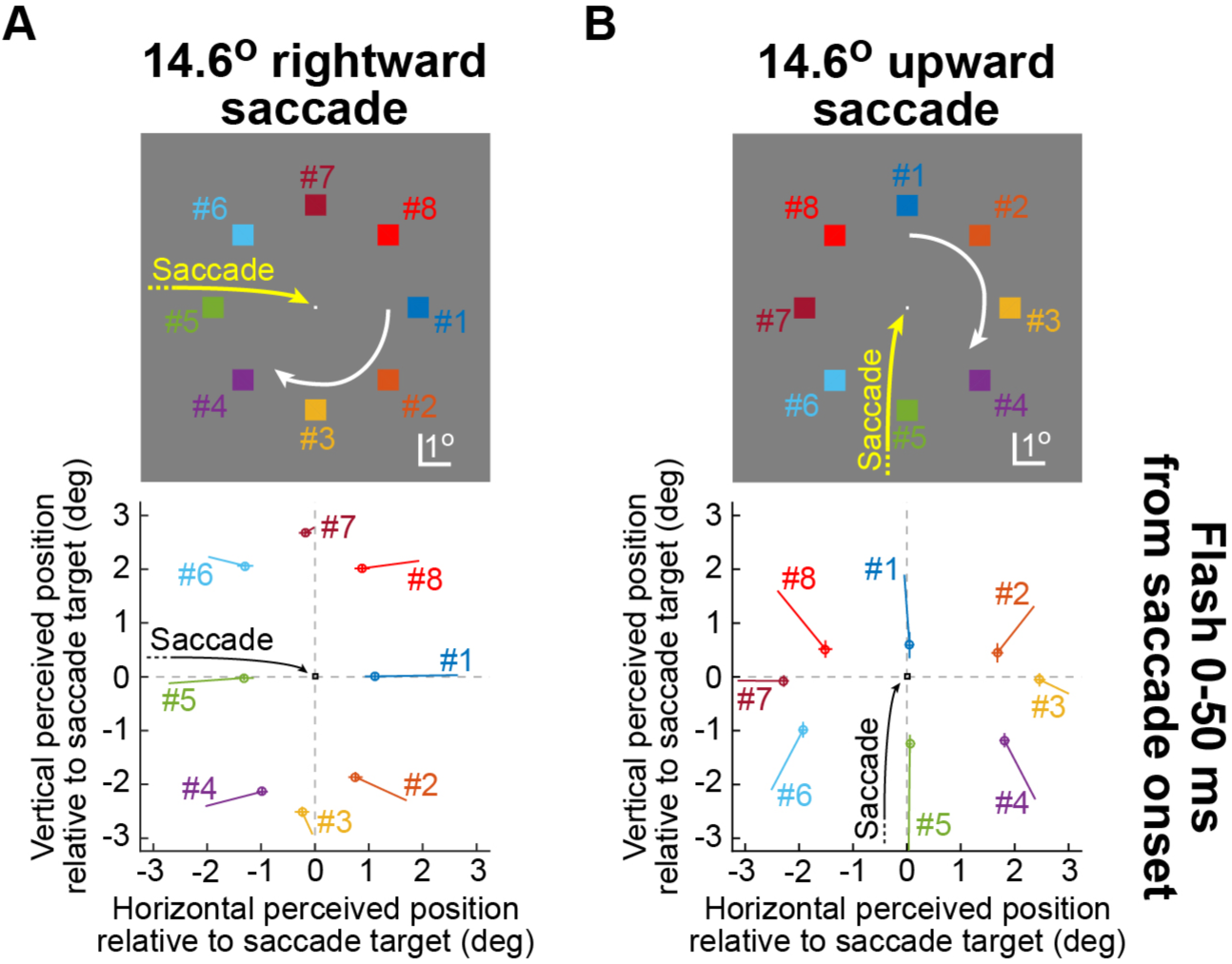
Patterns of peri-saccadic perceptual mislocalization for rightward versus upward saccades. (**A**) The top panel depicts our conventions for relating flash location to saccade direction. The yellow arrow indicates a rightward saccade towards the small white target presented in the middle of the schematic; the colored numbered squares indicate the possible peri-saccadic brief flash locations relative to the saccade target. For example, flash #1 indicates that a brief white flash appeared along the axis of the saccade direction and more eccentric than the saccade target (relative to initial fixation), and similarly for the other flashes (Materials and Methods). We numbered flash locations in a clockwise manner indicated by the white circular arrow. The bottom panel shows the perceived position of each flash location (relative to the saccade target location) for flashes occurring 0-50 ms from saccade onset. Error bars denote s.e.m. across trials. Each data point is depicted at the end of a line whose other endpoint indicates the perceived position of the same flash when it occurred 75-125 ms before saccade onset: peri-saccadic flashes were perceived as compressed towards the saccade target compared to the percept a long time before the saccade, as observed previously. N=53-73 trials per flash location in the interval 0-50 ms from saccade onset, and N=54-70 trials per flash location in the interval 75-125 ms before saccade onset. (**B**) The same analysis applied for upward saccades. The numbering conventions for flash locations relative to saccade target location and saccade direction are identical to **A**. The bottom panel shows that upward saccades were also associated with peri-saccadic compression, but that the compression effect was markedly stronger than in **A**. For example, flashes #1, #2, and #8 were associated with a closer perceived position to the saccade target than for rightward saccades (further illustrated and quantified in other figures). N=32-71 trials per flash location in the interval 0-50 ms from saccade onset, and N=61-80 trials per flash location in the interval 75-125 ms before saccade onset.

### Laboratory setup

The laboratory setup was similar to that described in (Bellet et al. 2017; Hafed 2013; Tian et al. 2016). Briefly, subjects sat in a dark room in front of a display running at 85 Hz and spanning 34.1 × 25.9 deg. Stimuli were white (97.3 cd/m^2^) and presented over a uniform gray background (20.5 cd/m^2^). Head fixation was achieved using a custom-made chin-and-forehead rest described previously (Hafed 2013), and we tracked eye movements using a video-based eye tracker running at 1 KHz (EyeLink 1000, SR Research). Subjects indicated their perceived flash location (see **Behavioral tasks** below) by pressing a computer mouse button after moving the mouse pointer to the desired point on the display. We used the Psychophysics Toolbox (Brainard 1997; Kleiner et al. 2007; Pelli 1997) for real-time display control.

### Behavioral tasks

In the main experiment with cardinal saccade directions (e.g. Fig. 1), trials started with the onset of a white fixation spot of ∼7.3×7.3 min arc dimensions. The spot appeared ∼7.3 deg from the center of the display in one of the four cardinal directions (right, left, up, or down relative to display center), and it was then jumped by ∼14.6 deg in the opposite direction after 1-3 seconds. That is, if the spot appeared ∼7.3 deg to the right of display center at trial onset, it jumped to ∼7.3 deg to the left of display center after 1-3 seconds, and similarly for the other initial fixation spot locations. In all cases, the fixation spot jump was the cue to generate a visually-guided saccade (∼14.6 deg in amplitude), as quickly as possible, from the old spot location to its new one. We started trials with the fixation spot being displaced from display center because we wanted to employ fairly large saccades (∼14.6 deg in amplitude) for all saccade directions, which would have been impossible for vertical saccades using our display’s aspect ratio. Using relatively large saccades was instrumental to allow us to compare our observations to those in classic investigations of peri-saccadic perceptual mislocalization, in which large saccades were also used (Cai et al. 1997; Kaiser and Lappe 2004; Ross et al. 1997; Ross et al. 2001).

In every trial, a white square of ∼0.73×0.73 deg dimensions was flashed for only one display frame (∼12 ms) after one of five different latencies from the fixation spot jump (∼60, 106, 153, 212, or 259 ms). Given that our observed saccadic reaction times were typically >100 ms and<300 ms, this meant that relative to saccade onset, the flash could come either before, during, or after an eye movement (with a relatively uniform distribution), allowing us to map time courses of peri-saccadic percepts. After a short delay from the flash presentation (500 ms), a mouse pointer was made visible always in the center of the display, and subjects were asked to click (within<3500 ms) on the location where they saw the flash. If subjects did not see any flash, they were instructed to click at the center of the display, but this event was rare (only 1.74% of accepted trials had a click near the center of the display). Subjects were otherwise encouraged to localize the flash accurately rather than making speeded clicks (manual reaction times were 1270 ms +/−360 ms s.d.; median: 1201 ms).

The flash could appear centered on one of 8 possible equally-spaced positions around a virtual circle surrounding the saccade target position (e.g. see Fig. 1). This virtual circle had a radius of ∼3.6 deg. In other words, if the fixation spot jumped to a new location, instructing a saccade, the flash could appear at an eccentricity of ∼3.6 deg from the new fixation spot location in one of 8 equally-spaced directions. Because prior experiments have revealed that two-dimensional patterns of peri-saccadic mislocalization depend on flash eccentricity relative to saccade target eccentricity (Kaiser and Lappe 2004), we always labeled flash positions during our analyses using a relative numbering convention, regardless of saccade direction. Specifically, flash position #1 was always the flash along the saccade direction and farther away from initial fixation than the saccade target.

Flashes #2-#8 were then the successive flashes along the virtual circle if one were to traverse this circle in a clockwise manner (e.g. see Fig. 1). Thus, using this convention, flashes #1, #2, and #8 were always the flashes that were more eccentric than the saccade target position relative to initial fixation position before the saccade, and flashes #4, #5, and #6 were always the flashes that were closer to initial fixation than the saccade target. Similarly, flashes #2, #4, #6, and #8 were always the flashes for which both “parallel” and “orthogonal” mislocalizations relative to the vector of the saccade (e.g. see Figs. 1, 5A) could occur (Kaiser and Lappe 2004).

In our second experiment, we had subjects perform 45-deg oblique saccades. The subjects made either rightward-upward or rightward-downward saccades. The start and end points of the fixation spot (relative to display center) were adjusted accordingly such that saccade amplitude was similar to the cardinal saccade direction experiment described above (∼14.6 deg). Also, the displacement of the fixation spot position from display center before and after the saccade was always symmetric relative to display center, exactly as in the first experiment above. Finally, flash times and locations relative to the saccade target location and saccade direction vector, as well as all flash numbering and naming conventions, were identical to those described above.

### Data analysis

We detected saccades using eye velocity and acceleration criteria (Chen and Hafed 2013), and we manually inspected all detected movements. We identified the saccade of interest (i.e. the movement responding to the jump in fixation spot location), and we analyzed its associated percepts. We excluded from analyses all trials in which a blink or microsaccade occurred between −200 ms from saccade target jump until the saccade was executed. We also excluded trials with saccade reaction times<100 ms or >500 ms from the jump in fixation spot, as well as trials with saccade endpoints being >4.5 deg from the saccade target (e.g. Fig. 4A, B), in order to ensure that subjects made appropriate target-directed movements. Because of variability in saccade metrics for upward versus downward saccades (Hafed and Chen 2016), in other analyses, we only included saccades with endpoints overlapping with endpoints of movements of saccades made in all other directions (e.g. Fig. 4). In other words, we only considered saccades with “overlapping” endpoints across all executed saccade directions. This allowed us to establish that changes in mislocalization patterns for, say, upward saccades (e.g. Figs. 1-3) were not explained by variability in the metrics of the saccades themselves when compared to other saccade directions. We also checked whether saccadic peak velocity could explain our results (Ostendorf et al. 2007). Finally, we excluded any trials with percept reports (i.e. mouse button press locations) within a square of +/−2.44 × +/−2.44 deg from display center or with percept (manual) reaction times >3500 ms. These were trials in which subjects likely did not see the flash at all, perhaps due to blinks and other lapses. In total, we had 12,060 acceptable trials from the first experiment, out of which 9,836 could be used in the more stringent overlapping endpoints control analyses. For the second experiment (with less conditions), the trial numbers were 2,424 and 1,924, respectively.

**Figure 2.**
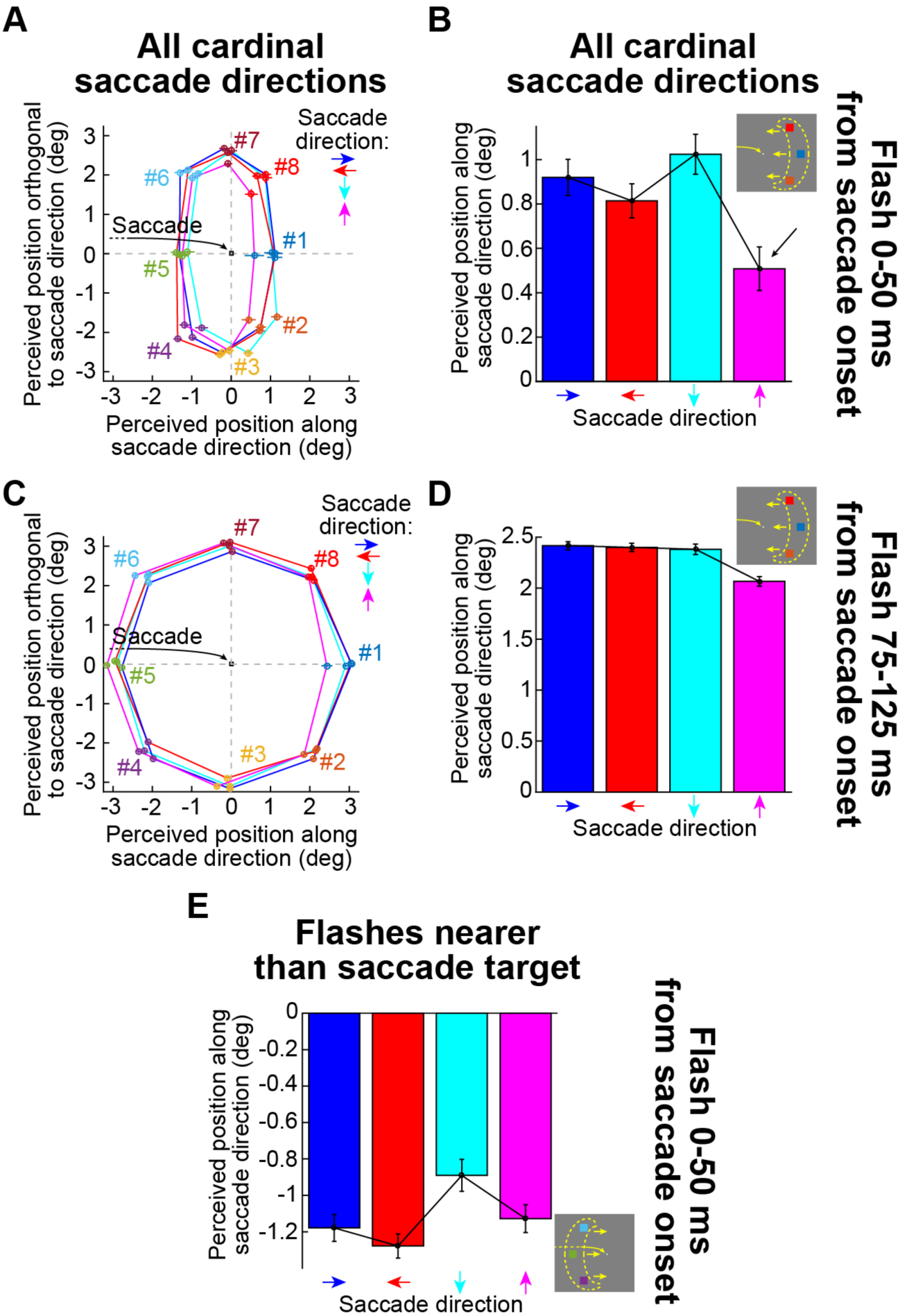
Peri-saccadic mislocalization was strongest for upward saccades with flash locations farther away from the saccade target location relative to initial fixation. (**A**) For flashes occurring 0-50 ms from saccade onset, we plotted the perceived position of each flash relative to the saccade target location (similar to Fig. 1), but now for each of the four cardinal saccade directions. We rotated the data such that the saccade vector in this figure was always rightward (despite the different saccade directions in the real experiment), but we maintained the relative positions of the flashes to the saccade vector in the rotation, such that flashes #1, #2, and #8 were always the flashes that were farther away from the saccade target, and flashes #4, #5, and #6 were always the flashes that were nearer than the saccade target (relative to initial fixation). Each solid line indicates the percepts associated with a given saccade direction according to the colored legend, and error bars denote s.e.m. The color coding of individual flash positions maintains the conventions set in Fig. 1. As can be seen, peri-saccadic percepts for upward saccades (magenta) were an outlier from all other saccade directions: flashes #1, #2, and #8 were significantly more compressed towards the saccade target. N=493, 493, 470, and 430 saccades for rightward, leftward, downward, and upward saccades, respectively (roughly equally divided across all flash locations). (**B**) For flashes #1, #2, and #8 (i.e. farther than the saccade target), we plotted the component of perceived flash position along the saccade direction (i.e. the x-axis values from **A**) for all four cardinal saccade directions. Upward saccades were associated with the strongest peri-saccadic compression towards the saccade target (i.e. the smallest values of perceived position relative to saccade target location; black arrow). Error bars denote s.e.m. (**C**, **D**) Similar analyses for flash times longer after saccade onset, demonstrating that the stronger mislocalization for upward saccades lingers for a longer time than in all other saccade directions. N=547, 576, 441, and 459 saccades for rightward, leftward, downward, and upward saccades, respectively (roughly equally divided across all flash locations). (**E**) An analysis similar to **B** but for all flashes nearer to initial fixation than the saccade target (flashes #4, #5, and #6; see schematic in the inset). The y-axis shows perceived position of the flashes (relative to the saccade target location) along saccade direction for all flashes occurring 0-50 ms from saccade onset. Unlike for flashes farther away from fixation (**B**), upward saccades showed similar levels of compression along the saccade direction vector as all other cardinal saccade directions (i.e. upward saccades were not an outlier). N=190, 177, 180, 171 trials for rightward, leftward, downward, and upward saccades, respectively.

**Figure 3.**
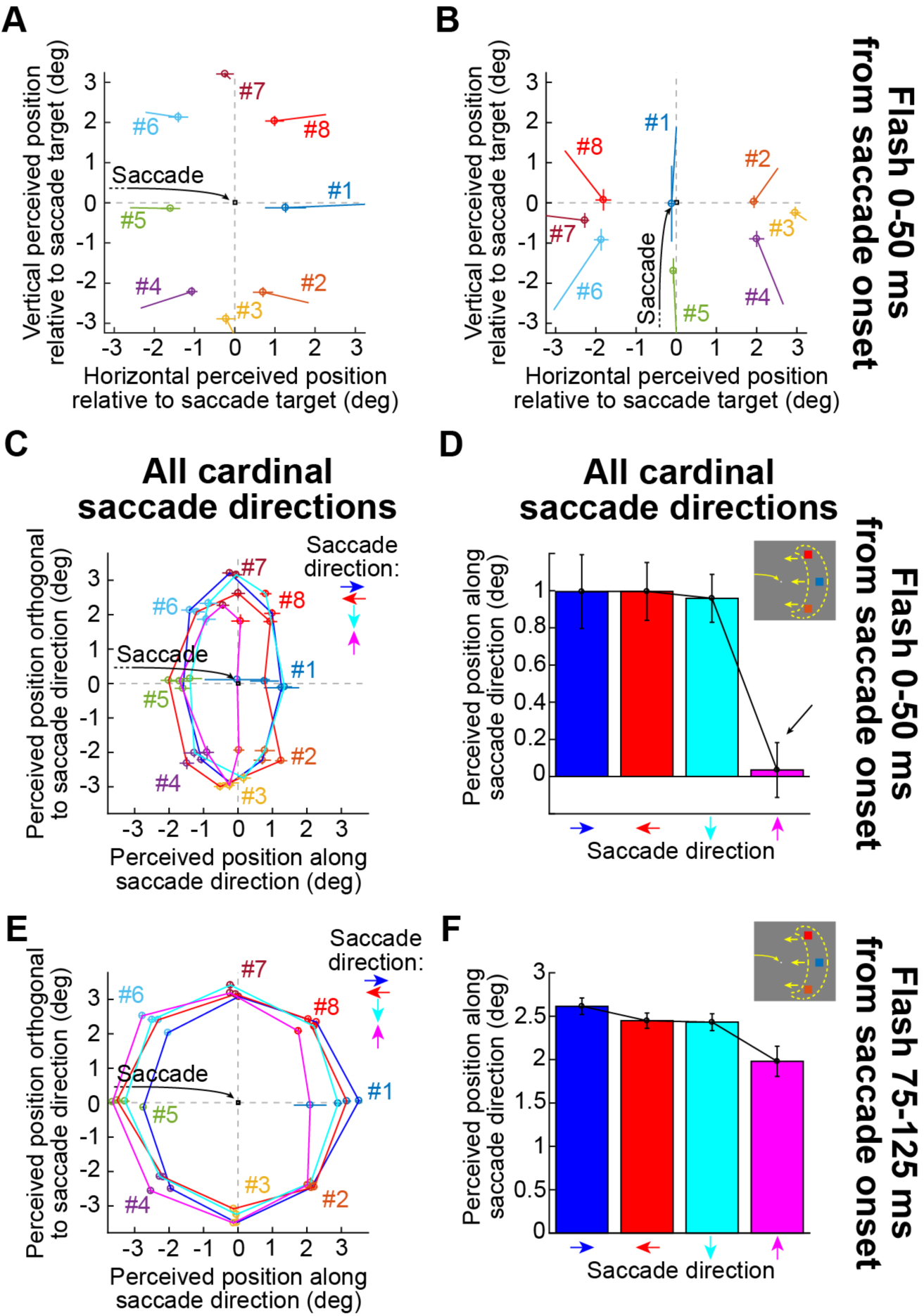
Peri-saccadic mislocalization patterns from a single exemplary subject. This figure is formatted identically to Figs. 1, 2, but using data from an example subject. The subject exhibited the same patterns of results as those seen in the aggregate analyses of Figs. 1, 2. (**A**) N=10-18 trials per flash location for the time interval 0-50 ms from saccade onset, and N=11-19 trials per flash location for the time interval 75-125 ms before saccade onset. (**B**) N=3-19 trials per flash location for 0-50 ms from saccade onset, and N=11-21 for 75-125 ms before saccade onset. (**C**, **D**) N=113, 125, 125, and 101 saccades for rightward, leftward, downward, and upward saccades, respectively. For **D**, p=1.252×10^−4^, 1-way ANOVA. (**E**, **F**) N=141, 136, 104, and 90 saccades for rightward, leftward, downward, and upward saccades, respectively. For **F**, p=0.0016, 1-way ANOVA.

**Figure 4.**
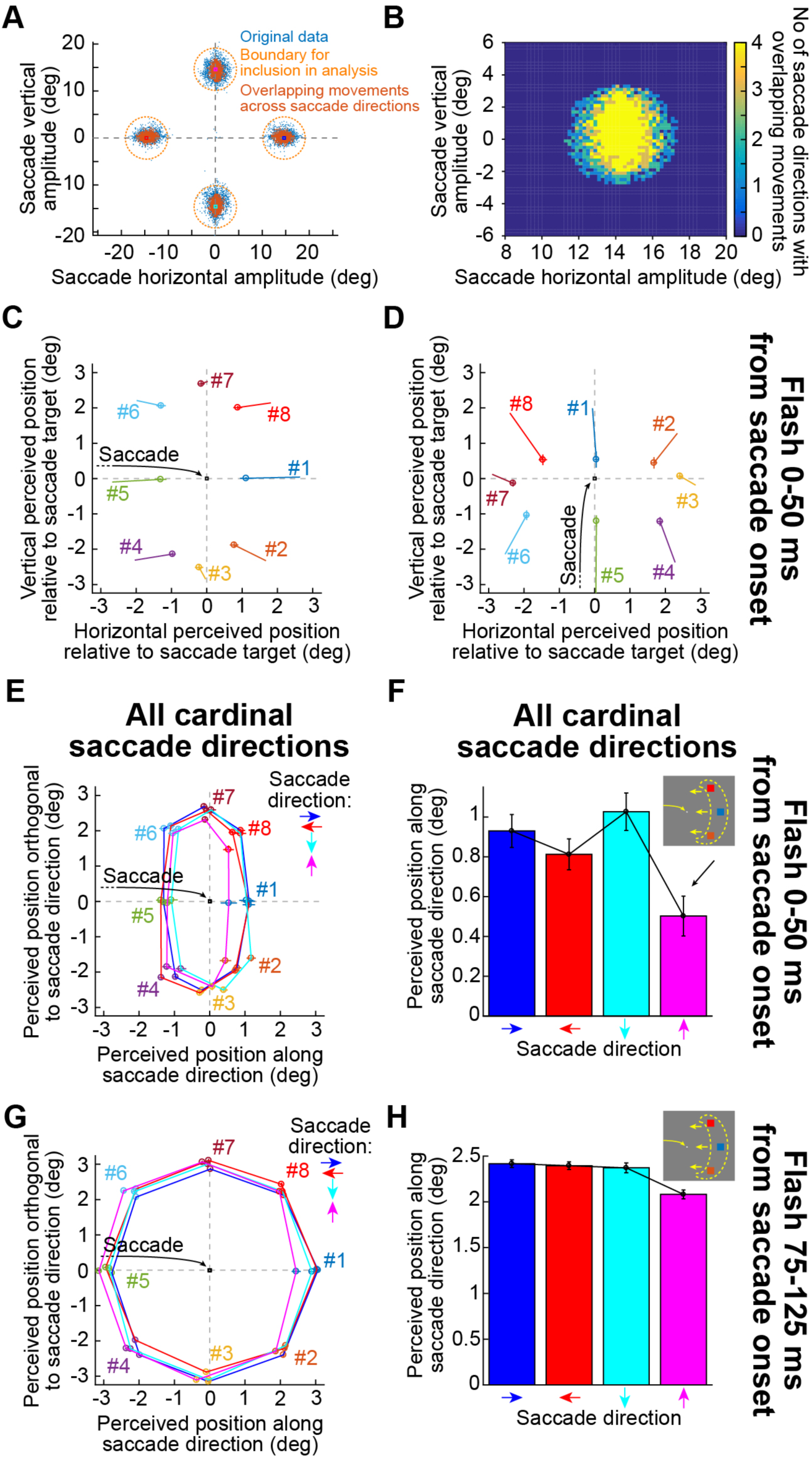
Controlling for variability of saccade endpoints. (**A**) Each blue dot shows the horizontal and vertical amplitude of a saccade; colored squares show the saccade target positions. We only analyzed trials in which saccades landed within the dashed orange circle for each saccade direction to ensure that subjects made a proper saccade upon instruction to do so. We also excluded saccadic and/or manual reaction time outliers in all analyses (Materials and Methods). In additional control analyses, we only analyzed trials in which saccade amplitudes and directions had overlap across all analyzed trials among the four saccade directions in this experiment (brown dots). In other words, if all the brown dots were rotated to aligned with a rightward saccade direction, then all four saccade directions would have matched variability in saccadic endpoints (**B**) We established the regions of overlap in **A** by rotating all saccade directions and plotting a cumulative histogram of saccade landing positions. The yellow region shows that this region was covered by all four saccade directions, and it was used to pick only the overlapping saccades of **A**. We used similar overlap criteria to also perform control analyses in our second experiment involving diagonal saccades. (**C**, **D**) Same analyses as in Fig. 1, but with only overlapping saccades. Stronger compression of flashes farther away from the saccade target relative to initial fixation in upward saccades still occurred. (**E**, **F**) Same analyses as in Fig. 2A, B, again showing that the results associated with upward saccades did not depend on potential differences in variability of saccadic landing positions across different saccade directions. For **F**, p=6.604×10^−4^, 1-way ANOVA. (**G**, **H**) Similar analysis at a different time point, again replicating the results of Fig. 2.

We measured perceived flash location as a function of saccade direction and flash time relative to saccade onset. We defined a peri-saccadic interval of interest in primary analyses as the interval in which flash times occurred 0-50 ms after saccade onset, because this is typically when maximal mislocalization is expected to occur, but we also saw mislocalization in other intervals (e.g. Fig. 5). We also compared this interval of interest to a pre-saccadic baseline interval with flash times occurring between −125 ms and −75 ms relative to saccade onset. To obtain time courses of percepts, we binned flash times relative to saccade onset in 50-ms bins, and we stepped the binning windows in steps of 2 ms. To display a spatial trajectory of perceived flash locations, we plotted all time course points in an interval around saccade onset (e.g. −100 ms to +100 ms) as spatial coordinates (e.g. vertical perceived position versus horizontal perceived position). The trajectories typically started from a baseline perceived position and then moved closer towards the saccade target location and then relaxed back to near the original perceived position (e.g. Figs. 5A, 7A, B, 8D). Therefore, the trajectories of percepts made a loop in position space. We estimated the direction of mislocalization by creating a vector whose origin was the midpoint of percepts at −100 ms or +100 ms relative to saccade onset (i.e. the baseline percept), and whose endpoint was either the point of maximal orthogonal or parallel mislocalization (depending on the specific goal of a given analysis) between −100 ms and +100 ms from saccade onset (typically occurring somewhere in the interval from 0 to 50 ms after saccade onset).

**Figure 5.**
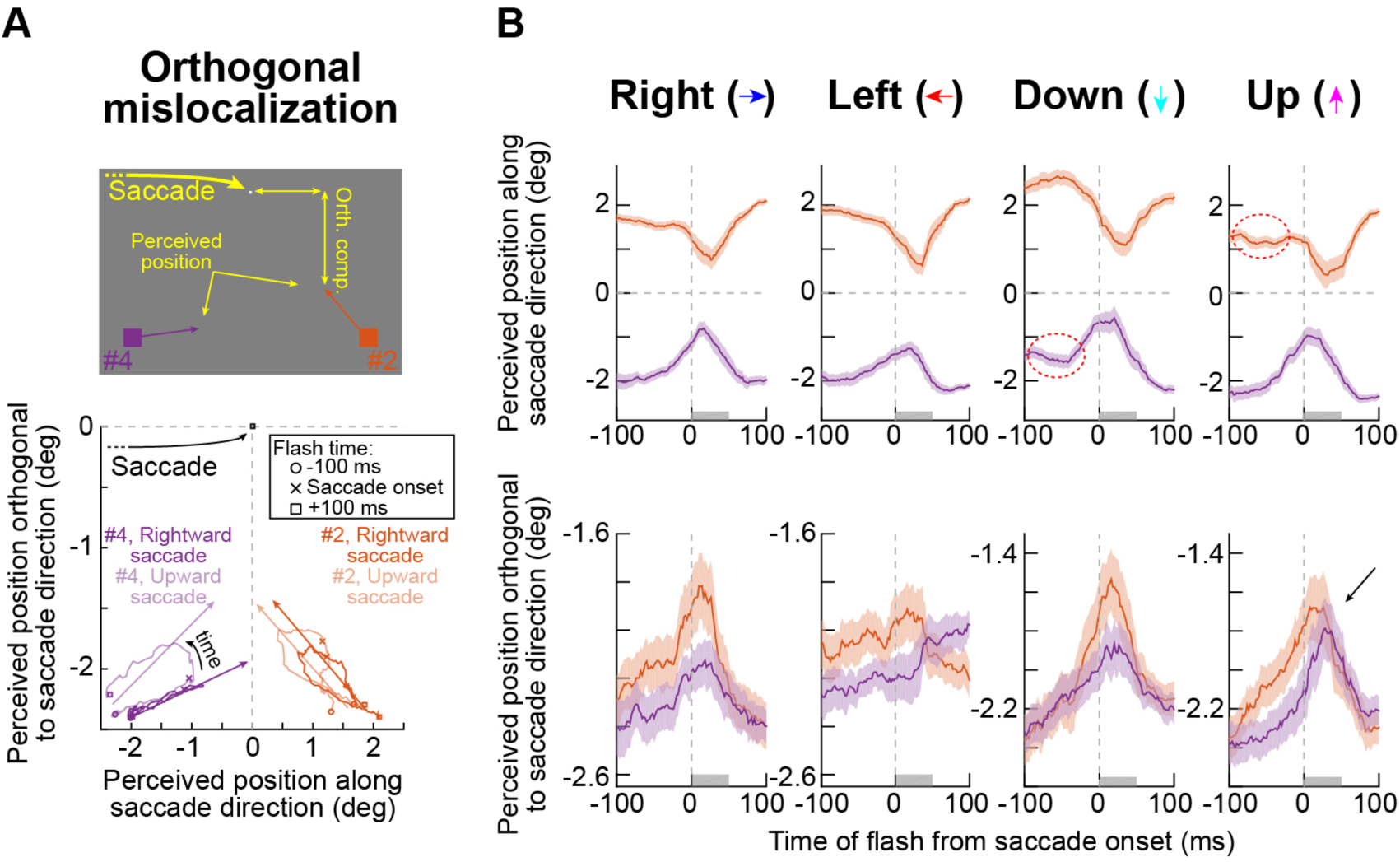
Mislocalization orthogonal to saccade direction was strongest for upward saccades, even for flash locations nearer to initial fixation than the saccade target. (**A**) It was previously shown that mislocalization orthogonal to saccade direction is asymmetric for flashes nearer or farther away from the saccade target. For example, the top schematic shows a rightward saccade and flashes #2 and #4. The orthogonal perceived position of flash #2 (“Orth. comp.”) should be closer to the saccade target than that of flash #4, resulting in “mislocalization vectors” having different slopes (orangish and purplish arrows). In the bottom panel, we plotted the trajectory of perceived position for these two flash locations from −100 ms to +100 ms relative to saccade onset, and we did this for either rightward or upward saccades (rotated as in Figs. 2-4 to facilitate comparison to rightward saccades). For rightward saccades (saturated colors), we confirmed the difference in mislocalization vector slopes: for each flash, the arrow with the saturated color connects the midpoint of the percepts at −100 ms and +100 ms with the position in the percept trajectory having maximal orthogonal component; the mislocalization vector for flash #4 had shallower slope than for flash #2. For upward saccades (unsaturated colors), orthogonal mislocalization was equally strong between nearer and farther flash locations (compare the milsocalization vector slopes). N=359, 364 trials for flash locations #2 and #4, respectively, for the rightward saccade; N=342, 329 trials for the same flash locations for the upward saccade. (**B**) Time courses of perceived flash position (relative to saccade target location) as a function of time from saccade onset for different saccade directions (columns). The top row shows perceived position along saccade direction (error bars: 95% confidence intervals) demonstrating expected compression. The bottom row shows the component of perceived position orthogonal to saccade direction. For all but the upward saccade, during the interval 0-50 ms (shaded gray bars), orthogonal mislocalization was stronger (i.e. closer to zero y-axis values) for farther (flash #2, orangish) than nearer (flash #4, purplish) flashes. Upward saccades violated this observation (black arrow). Each time bin shown had N=47-98 repetitions. Other details in this figure, like the dashed red ellipses, are discussed later in the text.

**Figure 6.**
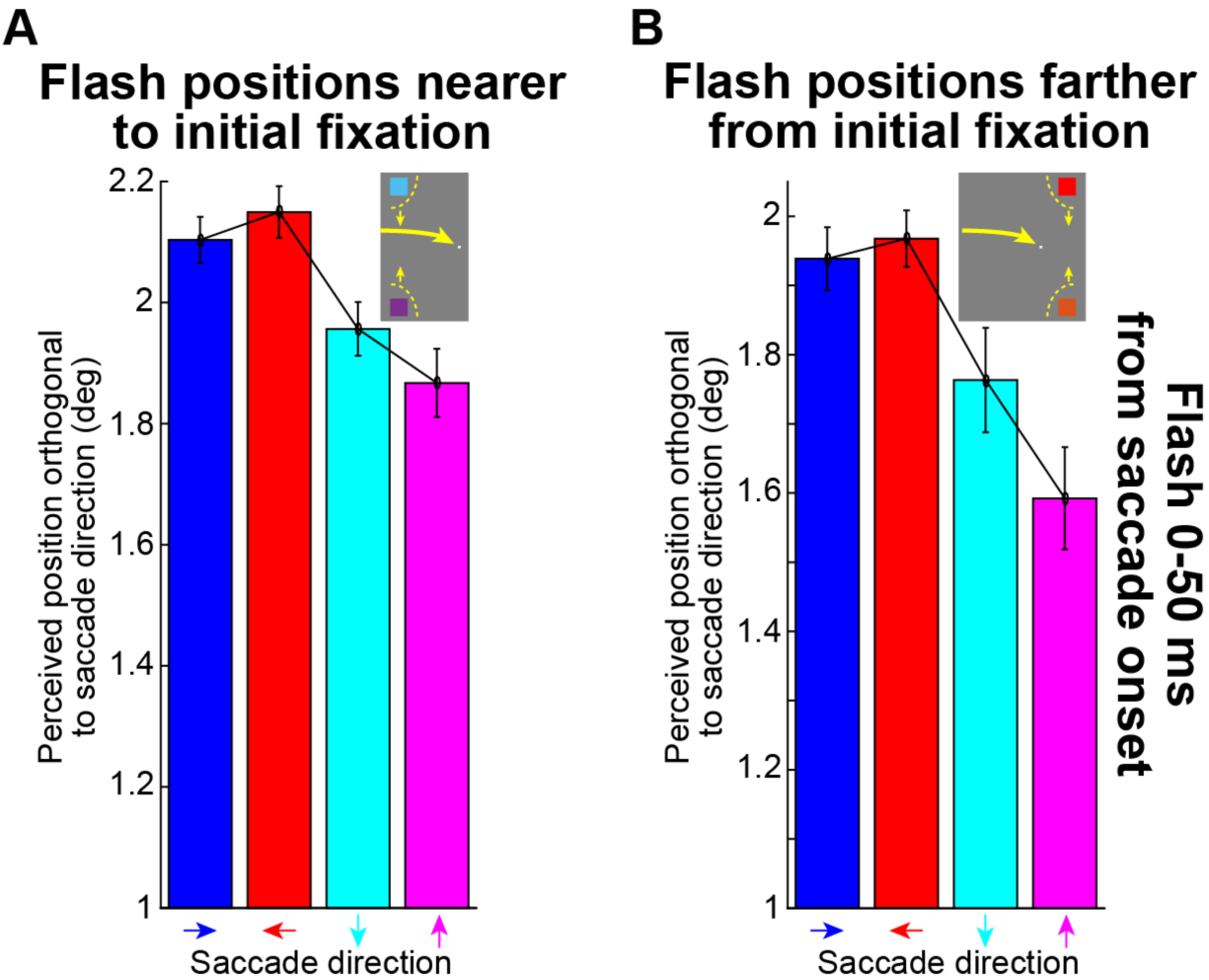
Orthogonal mislocalization relative to saccade direction was strong for upward saccades even for near flash locations. (**A**) We took all flash locations nearer to initial fixation than the saccade target, and that could still experience orthogonal mislocalization relative to saccade direction (i.e. flashes #4 and #6), and we plotted the average perceived position of the flash orthogonal to the axis of a saccade vector for flashes 0-50 ms from saccade onset. As in all analyses, perceived position was calculated relative to the saccade target location, meaning that smaller values in the plot indicate stronger compression. Consistent with Fig. 5, the upward saccade had the strongest orthogonal mislocalization even though the flashes analyzed were nearer to initial fixation than the saccade target. (**B**) This effect was magnified even more for all flashes farther away from the saccade target (i.e. flashes #2 and #8). Thus, orthogonal mislocalization was prominent for upward saccades, even for flashes nearer to initial fixation than the saccade target. Note that the y-axes are different in the two panels, and they confirm that orthogonal mislocalization for horizontal saccades is stronger for farther than nearer flash locations. Error bars denote s.e.m. N=126, 115, 124, and 123 trials for rightward, leftward, downward, and upward saccades, respectively in **A** and N=116, 126, 20, and 97 trials for rightward, leftward, downward, and upward saccades in **B**.

**Figure 7.**
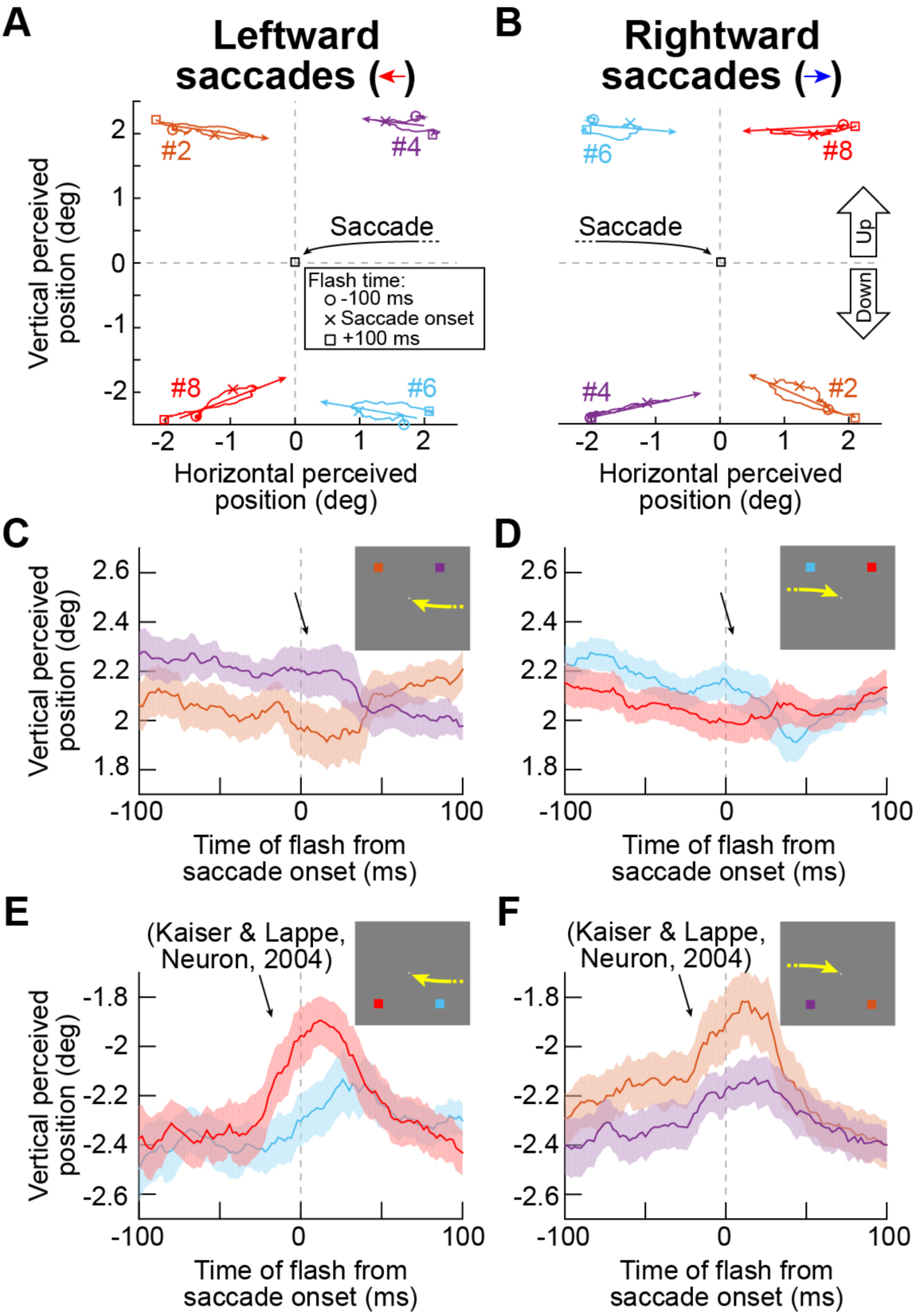
For horizontal saccades, mislocalization patterns differed between flashes in the upper versus lower retinotopic visual fields. (**A**, **B**) We took a closer look at the phenomenon of orthogonal mislocalization from Figs. 5, 6. For leftward (**A**) or rightward (**B**) saccades, we plotted mislocalization trajectories (similar to Fig. 5A) for all flash locations that were either farther (flashes #2 and #8) or nearer (flashes #4 and #6) from initial fixation than the saccade target, and that could still experience orthogonal mislocalization. Each curve plots the percept trajectory for a given flash location from −100 ms to +100 ms from saccade onset, and the arrow connects the baseline percept (midpoint of −100 ms and +100 ms percepts) to the point in each percept trajectory with maximal horizontal mislocalization. Independently of saccade direction, flashes in the lower visual field (flashes #6 and #8 for leftward saccades and flashes #4 and #6 for rightward saccades) had asymmetric orthogonal mislocalization (differences in mislocalization vector slopes between nearer and farther flash locations), as we saw for the rightward saccade in Fig. 5, and as also seen previously. However, flashes in the upper visual field only experienced parallel mislocalization with no orthogonal component (mostly horizontal mislocalization vectors). (**C**, **D**) This observation was confirmed when we plotted time courses of vertical perceived position of the upper visual field flashes. There was no strong peri-saccadic modulation in vertical perceived position (diagonal arrows). (**E**, **F**) However, for flashes in the lower visual field, vertical perceived position changed peri-saccadically, and the mislocalization patterns also confirmed previous observations that farther flash locations (flash #8 for leftward saccades and flash #2 for rightward saccades) experienced stronger orthogonal mislocalization than nearer flash locations (flash #6 for leftward saccades and flash #4 for rightward saccades). Thus, retinotopic upper or lower visual field flash location is a contributor to differences in percepts for different saccade directions. Error bars denote 95% confidence intervals, and each time bin included 50-101 repetitions.

**Figure 8.**
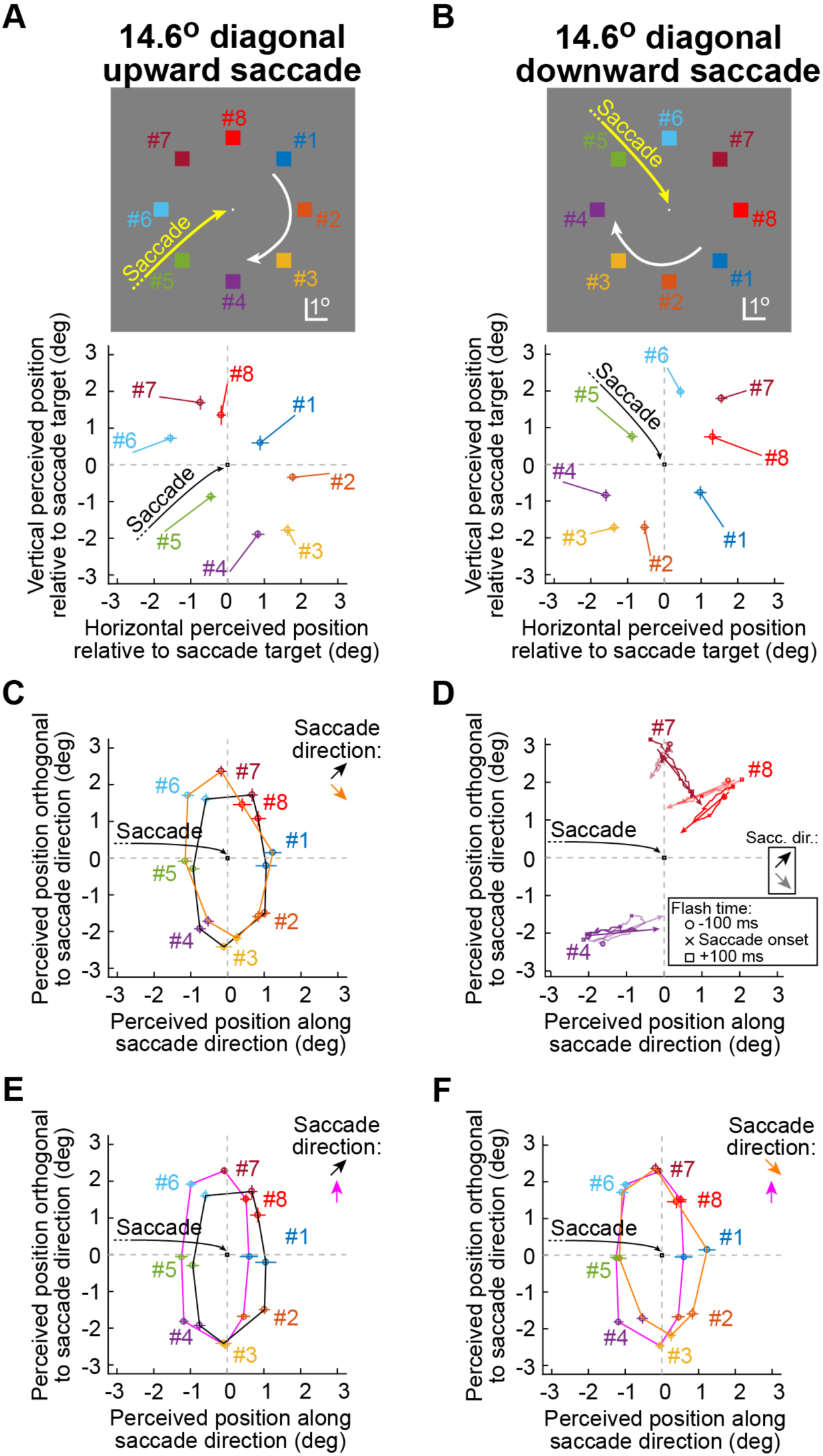
Diagonal saccades revealed mislocalization asymmetries beyond classic compression based on flashes being farther or nearer from the saccade target relative to initial fixation. (**A**, **B**) We plotted perceived flash position (for flashes occurring 0-50 ms from saccade onset) for either diagonal upward (**A**) or diagonal downward (**B**) saccades. The top schematics show identical flash locations relative to the saccade target as in our earlier experiment, and with similar conventions relative to saccade target location and saccade direction. The bottom panels show percept locations, along with s.e.m. error bars. The origin of the line connected to each individual data point illustrates the location of the percept when the corresponding flash occurred 75-125 ms before saccade onset (as in Fig. 1). Both diagonal saccades were associated with compression towards the saccade target, but there were notable differences in the patterns of compression. (**C**) We rotated both saccades to our standard reference frame to demonstrate the differences in compression between diagonal upward and diagonal downward saccades. Diagonal upward saccades showed substantially stronger compression for flash locations (like #7 and #8) that were located physically higher in the display relative to the saccade target than for diagonal downward saccades. Figure 9 shows the same analysis as here but using data from a single subject to demonstrate robustness of the phenomenon even on the single-subject level. (**D**) Percept trajectories (as in Figs. 5, 7) for example flash locations demonstrating similarities (flash #4) or differences (flashes #7 and #8) between the diagonal upward (saturated colors) and diagonal downward (unsaturated colors) saccades. This figure is formatted identically to a similar analysis in Fig. 7. Supplementary Movie 1 shows an animation of these trajectories. N=180 and 198 trials for diagonal upward and diagonal downward saccades, respectively, in the interval 0-50 ms from saccade onset (**A**-**C**). Also, each time bin in the trajectories of **D** had 15-44 repetitions. (**E**) Data from the diagonal upward saccade (black) plotted along with those from a purely upward saccade (magenta) for comparison. Diagonal upward saccades were associated with bigger distortions in percepts for flashes nearer to initial fixation than the saccade target. (**F**) Same comparison to upward saccades but for the diagonal downward saccades (orange). Diagonal downward saccades were associated with different patterns of percepts than the diagonal upward saccades. Thus, patterns of peri-saccadic mislocalization depend on saccade direction.

To compare perceptual mislocalization for different saccade directions, we rotated all data to align with a rightward saccade. For example, for an upward saccade, we rotated all data 90 degrees clockwise to facilitate comparison to rightward saccades. All flash locations relative to the saccade target and initial fixation position were consistently rotated as in the scheme of Fig. 1. This was critical to maintain relative eccentricity relationships for all flash locations across saccade directions, especially because it was previously suggested that it is relative eccentricity that seems to matter the most for patterns of two-dimensional peri-saccadic mislocalization (Kaiser and Lappe 2004; VanRullen 2004). Thus, we reported two-dimensional percept position in terms of position “parallel” to the saccade direction vector and position “orthogonal” to the saccade direction vector (e.g. Fig. 5A).

All figures in this paper report the numbers of trials used for the analyses, as well as measures of variability in the form of either s.e.m. error bars or 95% confidence intervals. Because peri-saccadic mislocalization is a highly robust phenomenon (Ross et al. 1997), with many replications in the field over at least two decades, we pooled data from all subjects for most of our analyses. However, we confirmed that all of our observations can also be made at the individual subject level (e.g. Figs. 3, 9).

**Figure 9.**
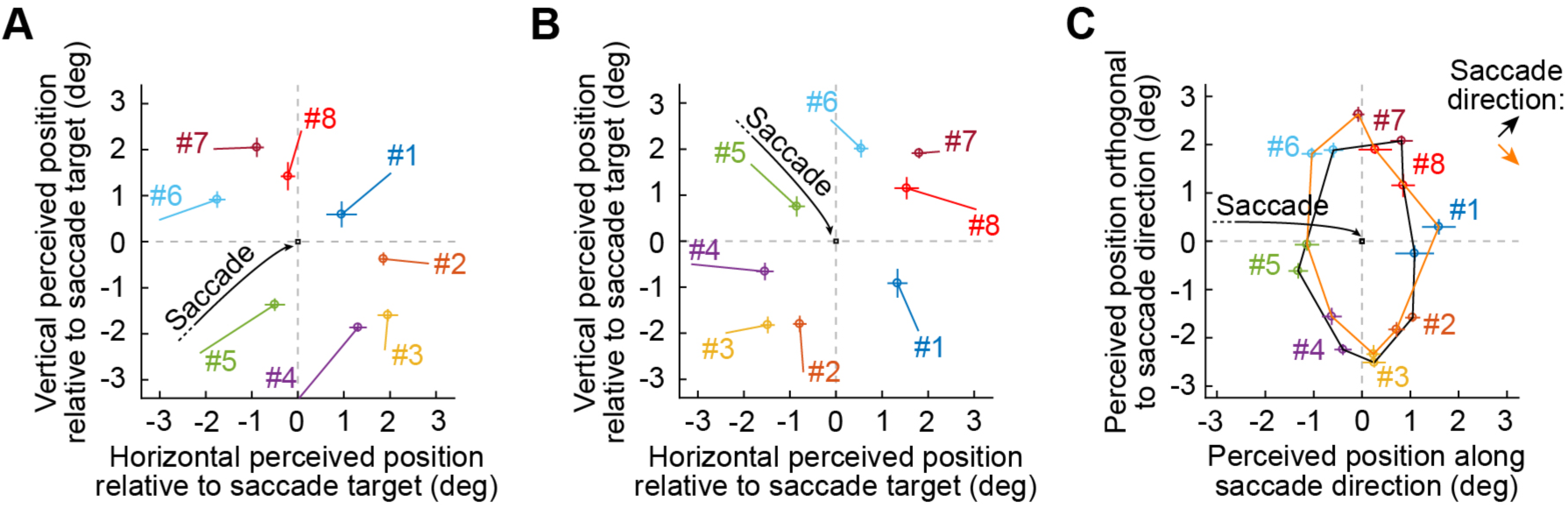
Peri-saccadic mislocalization for diagonal saccades in an exemplary individual subject. (**A**-**C**) This figure shows identical analyses to those shown in Fig. 8A-C but using data from only a single subject (same as in Fig. 3). The same differences between diagonal upward and diagonal downward saccades were evident even on an individual subject basis. Error bars denote s.e.m.

Finally, for conceptual modeling, we asked whether a simple mechanism of saccade-induced translation of visual location of stimuli, but in neural tissue coordinates, can account for perceptual mislocalization (VanRullen 2004). The specific question that we asked was on whether possible asymmetries in tissue representations between the upper and lower visual fields are sufficient to make upward saccades look like outliers in terms of perceptual mislocalization. As such, we modeled visuo-motor maps as having both foveal magnification, as in (VanRullen 2004), as well as upper versus lower visual field asymmetries, as in (Hafed and Chen 2016). The equations for the maps with upper and lower visual field asymmetries were identical to those in (Hafed and Chen 2016). The equations for perfectly symmetric maps were similar, but without the area scale factor invoked in (Hafed and Chen 2016). Note that these maps were based on superior colliculus (SC) topography, but their mathematical form is identical to other visual maps in the brain (e.g. primary visual cortex, V1). Specifically, the maps employ logarithmic warping such that foveal visual image locations are associated with a greater area of neural tissue. Naturally, maps mathematically fit to the size of V1 or other visual or motor areas can be potentially used, but our goal here was to conceptually ask whether asymmetries in maps between upper and lower visual field representations can be related to our experimental results. Thus, specific map parameters (fit to V1 or SC or elsewhere) are less critical than the idea that a map may or may not have upper and lower visual field asymmetries.

## Results

We ran four human participants on an experiment involving brief stimulus flashes presented around the time of either rightward, leftward, downward, or upward saccades. Subjects localized the flashes at the ends of trials by manually pointing a cursor to where they perceived them. Flashes could occur from approximately −100 ms to approximately +100 ms from saccade onset, and with roughly uniform probability (Materials and Methods).

We first replicated observations of two-dimensional peri-saccadic perceptual mislocalization for rightward saccades (Kaiser and Lappe 2004). In our experiment, a brief flash could appear at any one of eight possible locations lying on a virtual circle surrounding the saccade target (Fig. 1A, top). Because mislocalization patterns depend on flash eccentricity from initial fixation position relative to saccade target eccentricity (Kaiser and Lappe 2004; VanRullen 2004), we classified flashes as being either more (“farther”) or less (“nearer”) eccentric than the saccade target (relative to initial fixation), and using the numbering convention shown in Fig. 1A (top). Thus, flashes #1, #2, and #8 were all farther from initial fixation than the saccade target, and were therefore expected to be misperceived backwards (Kaiser and Lappe 2004); flashes #4, #5, and #6 were nearer to initial fixation, and were expected to be misperceived forwards (Kaiser and Lappe 2004). The net result is a “compression” of percepts towards the saccade target as shown in Fig. 1A (bottom). In this figure, each data point indicates average perceived location for flashes presented 0-50 ms from saccade onset, and each line connects each data point to its corresponding percept when flashes were presented ∼100 ms before saccade onset (Materials and Methods) (i.e. when perception was expected to be more veridical). Two-dimensional compression of perceived flash location took place (Kaiser and Lappe 2004).

For upward saccades (Fig. 1B), notable differences were evident to us and are the primary focus here. For example, using the same conventions as in Fig. 1A, flash #1 for upward saccades (Fig. 1B, top) was more eccentric than the saccade target relative to initial fixation position (Fig. 1A, top). However, the peri-saccadic percept associated with this flash location was closer to the saccade target than for rightward saccades (Fig. 1B, bottom). In other words, at the time of expected peak mislocalization relative to saccade onset (0-50 ms from saccade onset), mislocalization for flash #1 (and #2 and #8) was stronger for upward saccades than for rightward ones (compare Figs. 1A and 1B). In what follows, we quantify this and other differences in two-dimensional peri-saccadic perceptual mislocalization patterns between upward saccades and other cardinal saccade directions, and we then explore potential origins and consequences of such differences.

### Stronger compression of farther flash locations for upward saccades

Using the same numbering convention of flash locations as in Fig. 1, we compared peri-saccadic perceptual mislocalization patterns for different saccade directions using a single reference frame (namely, flash eccentricity relative to saccade target eccentricity). This is because accounts of peri-saccadic mislocalization critically rely on relative eccentricities of flashes and saccade targets (Hamker et al. 2008; Hamker et al. 2011; Kaiser and Lappe 2004; Richard et al. 2009; VanRullen 2004; Zirnsak et al. 2010). In Fig. 2A, we plotted perceived flash location (relative to the saccade target) for all flashes occurring 0-50 ms from saccade onset, and we connected all data points from all eight flash locations for a given saccade direction with a contour having a single color (four colors denote the four saccade directions). To facilitate comparison of peri-saccadic mislocalization for all saccade directions, we rotated all data points to align with the rightward saccade, and flash locations were also rotated to respect the conventions of Fig. 1 (Materials and Methods). For example, regardless of saccade direction, flash #1 was more eccentric than the saccade target and along the vector of the saccade (“parallel” to the saccade vector), and flash #5 was less eccentric and similarly along the vector of the saccade; on the other hand, flash #2 was more eccentric than the saccade target but at a clockwise-shifted angle relative to flash #1 (i.e. having a component of position “orthogonal” to the saccade vector), and so on for all other flash locations. If perception was veridical, then the shown data points should all lie on a circle (e.g. Fig. 1A, top). However, percepts were compressed towards the saccade target. Critically, percepts for upward saccades were outliers when compared to all other saccade directions: perceived flash position along the saccade direction vector for flashes #1, #2, and #8 was closer to the saccade target location (i.e. closer to zero in the x-axis of the figure) for upward saccades than for all other saccade directions. This means that there was stronger compression along the saccade direction for flashes farther away from the initial fixation position than the saccade target.

To summarize this observation, we plotted average perceived position parallel to the saccade vector for flashes #1, #2, and #8 (i.e. all flashes farther way from initial fixation than the saccade target). For flashes 0-50 ms after saccade onset (Fig. 2B), such perceived position was similar for rightward, leftward, and downward saccades, but upward saccades were associated with stronger compression (i.e. smaller values of perceived position along the saccade vector) (p=6.295×10^−4^, 1-way ANOVA). Thus, backwards compression of flashes farther than the saccade target was stronger in peri-saccadic intervals for upward saccades than for the other three saccade directions.

A longer time after saccade onset (e.g. ∼100 ms), percepts of flash location were more veridical, as expected (Fig. 2C), but the outlying nature of upward saccades still persisted even at such longer times (Fig. 2D) (p=1.672×10^−8^, 1-way ANOVA). This outlying nature of upward saccades was also specific for flashes more eccentric than the saccade target. When we repeated the same analysis of Fig. 2B but for flashes #4, #5, and #6 (i.e. nearer to initial fixation than the saccade target), peri-saccadic percepts for upward saccades appeared similar to those for the other saccade directions (Fig. 2E; also see the nearer flash locations shown in Fig. 2a).

The above observations were also robust at the individual subject level. For example, Fig. 3 shows the same analyses as those made in Figs. 1-2 but for only one exemplary subject. All of the salient features of Figs. 1-2 can be seen. This is not surprising because of the robustness and replicability of peri-saccadic mislocalization in general (Ross et al. 1997).

We also wondered whether variability in saccade metrics might account for differences in peri-saccadic perceptual mislocalization for upward saccades. For example, saccade landing errors could be different for upward versus downward saccades (Hafed and Chen 2016; Schlykowa et al. 1996; Zhou and King 2002). We thus performed control analyses in which we only included data for analysis that had identical variability of saccade endpoints (Fig. 4A, B and Materials and Methods); in other words, there was maximal overlap in the distribution of saccade endpoints for all 4 saccade directions in these control analyses. We still replicated the findings of Figs. 1-2 (Fig. 4C-H). This means that the stronger backwards compression of farther flash locations for upward saccades was not an artifact of potential differences in saccade landing variability. We also checked whether peak velocity for upward saccades was inherently different from, say, rightward saccades, since prior work showed that compression strength does change with peak velocity (Ostendorf et al. 2007). This was not the case. For example, in the same data of Fig. 4, peak velocity for upward and, say, rightward saccades was not statistically different (p=0.053, t-test, N=2685 rightward saccades, N=2168 upward saccades).

Finally, our experimental design ruled out the possibility that mislocalization, and differences for upward saccades, were a result of a general bias to manually click near the center of the display, perhaps due to increased uncertainty (Brenner et al. 2008; Maij et al. 2010; Maij et al. 2011) about spatial and temporal reference frames peri-saccadically (Materials and Methods). Therefore, stronger peri-saccadic compression of farther flash locations for upward saccades was a robust observation.

### Stronger orthogonal mislocalization for upward saccades, regardless of flash eccentricity relative to the saccade target eccentricity

The above results have focused on misperceptions parallel to the saccade vector. However, some of our flashes were also off the axis of the saccade (e.g. flashes #2 and #4). Previous studies have shown that such flashes are additionally mislocalized along a direction orthogonal to saccades (i.e. overall, there is oblique perceptual mislocalization) (Kaiser and Lappe 2004). Moreover, whether a flash is farther (e.g. flash #2) or nearer (e.g. flash #4) to the saccade target dictates an asymmetry in orthogonal mislocalization (Kaiser and Lappe 2004), which current models account for by the fact that nearer eccentricities are magnified in neural tissue when compared to farther eccentricities (Hamker et al. 2008; Hamker et al. 2011; Kaiser and Lappe 2004; Richard et al. 2009; VanRullen 2004; Zirnsak et al. 2010).

Consider, for example, the scenario shown in Fig. 5A (top) for rightward saccades. Both flashes #2 and #4 would be expected to experience orthogonal mislocalization. However, orthogonal mislocalization is expected to be stronger for flash #2 than for flash #4 (Kaiser and Lappe 2004), such that if one were to draw a mislocalization “vector” (with origin being the percept long before or after a saccade and end being the percept peri-saccadically), then such a vector would be more oblique for flash #2 than for flash #4 (Fig. 5A, top). We replicated this finding (Kaiser and Lappe 2004) for rightward saccades (Fig. 5A, bottom, saturated colors); we plotted the trajectory of perceived flash position from −100 ms to +100 ms relative to saccade onset and in steps of 2 ms (Materials and Methods). We then created a vector whose origin was the midpoint of the percepts at - 100 and +100 ms relative to saccade onset (i.e. temporally far from saccade onset) and whose end was the point on the percept trajectory having maximal orthogonal mislocalization. As can be seen from the saturated colors in Fig. 5A (bottom), the mislocalization vector was indeed more oblique for flash #2 than for flash #4, even though the flashes were otherwise symmetric with respect to the saccade target (Kaiser and Lappe 2004). However, repeating the same analysis for upward saccades revealed a different pattern: orthogonal mislocalization was as strong for flash #4 as it was for flash #2 (Fig. 5A, bottom, unsaturated colors). Thus, for upward saccades, there was strong orthogonal mislocalization even for flashes nearer to initial fixation than the saccade target, unlike in (Kaiser and Lappe).

This observation was also evident when we plotted time courses of peri-saccadic perceptual mislocalization for flashes #2 and #4. In each column of Fig. 5B, the top panel illustrates perceived position (along with 95% confidence intervals) of either flash #2 or flash #4 along the dimension parallel to the saccade vector (Materials and Methods); the bottom panel illustrates perceived position (along with 95% confidence intervals) of the same flashes orthogonal to the saccade. Since the two flashes were either more or less eccentric than the saccade target, parallel perceived positions were expected to become compressed towards the saccade target (i.e. to move closer to zero in the top row) regardless of saccade direction. This was indeed the case (e.g. 0-50 ms from saccade onset; shaded gray bars in Fig. 5B). However, if neural maps responsible for peri-saccadic mislocalization were purely symmetric (Hamker et al. 2008; Hamker et al. 2011; Kaiser and Lappe 2004; Richard et al. 2009; VanRullen 2004; Zirnsak et al. 2010), then the orthogonal component of perceived position (Fig. 5B, bottom) should have revealed an asymmetry (Kaiser and Lappe 2004) like that predicted by Fig. 5A (top) for all saccade directions. This was violated for upward saccades (Fig. 5B, bottom row and diagonal black arrow in the rightmost panel); in the period of maximum mislocalization (shaded gray bars), orthogonal percepts were similar for flashes #2 and #4 with upward saccades, but not for the other saccade directions.

We next summarized these orthogonal mislocalization results, but for all off-axis flash locations that also had a nearer or farther eccentricity component relative to the saccade target (i.e. flashes #2, #4, #6, and #8). In Fig. 6A, we plotted the orthogonal component of perceived flash location for nearer off-axis flashes (i.e. flashes #4 and #6), and in Fig. 6B, we plotted the same variable for farther off-axis flashes (i.e. flashes #2 and #8). In both cases, orthogonal mislocalization clearly depended on saccade direction (p=3.02×10^−5^, 1-way ANOVA for Fig. 6A, and p=3.55×10^−5^, 1-way ANOVA for Fig. 6B).

Importantly, orthogonal mislocalization was always the strongest for upward saccades (i.e. the percept was closest to 0 in the figure; this is also evident in Fig. 2A and Fig. 3C). Therefore, upward saccades were not only associated with stronger parallel compression of farther flash locations (Figs. 1-4), but they were also associated with stronger orthogonal mislocalization even for nearer flash locations (Figs. 5-6).

We were also intrigued by several other details evident in the time courses of Fig. 5B. For example, for upward saccades, there was a biased percept in the parallel component of flash location for flash #2 long before saccade onset (Fig. 5B, top row, upward saccade, dashed red ellipse). This bias means that subjects reported flash #2 as being lower than it really was even 100 ms before saccade onset (well before the peri-saccadic transient deflection in percept started to occur). However, this does not necessarily mean that subjects had an overall bias to click slightly more downward only for upward saccades (or only for locations in the upper visual field relative to the head), and it also does not mean that our results above (Figs. 1-6) were fully explained by simple click biases associated with upward saccades. Specifically, the same early bias in percept well before saccade onset was also evident for downward saccades (i.e. with locations in the lower visual field relative to the head), but this time in association with flash #4 (Fig. 5B, top row, downward saccade, dashed red ellipse). In this case, subjects perceived this flash location as being more downward than it really was even 100 ms before saccade onset, and once again well before the peri-saccadic transient distortion in perception began to emerge. Interestingly, in both of these cases, flash #2 for upward saccades and flash #4 for downward saccades were in reality physically *above* the saccade target location. This means that in addition to peri-saccadic mislocalization being directly tied to saccade vectors (Kaiser and Lappe 2004; Ross et al. 1997), additional factors related to flash location itself (e.g. being physically up) matter. What is it, then, about *retinotopic* flash location that can influence percepts in our task? Related to this, why was the orthogonal percept in Fig. 5B (bottom) apparently so different in nature between rightward and leftward saccades, even though both were horizontal eye movements? We next turn to exploring these additional intriguing questions.

### Different mislocalization patterns of upper versus lower visual field flash locations for horizontal saccades

Even though our analyses of Fig. 6 summarized all off-axis flash locations having nearer or farther components (#2, #4, #6, and #8), Fig. 5B only showed time courses of mislocalization for flashes #2 and #4. In this figure, orthogonal mislocalization seemed to be conspicuously almost absent for the leftward saccade, and even for the flash #2 location (Fig. 5B, bottom, leftward saccade). This is very different from rightward saccades (Fig. 5A, saturated colors) (Kaiser and Lappe 2004). We explored this apparent discrepancy by comparing retinotopic flash locations in more detail. Specifically, because of our convention to relate flash eccentricity to saccade target eccentricity (Hamker et al. 2008; Hamker et al. 2011; Kaiser and Lappe 2004; Richard et al. 2009; VanRullen 2004; Zirnsak et al. 2010) independently of saccade direction, and because we rotated data to a single reference frame (Figs. 2-6), flashes #2 and #4 for the leftward saccade were in reality in the upper visual field retinotopically, whereas they were in the lower visual field for rightward saccades. So, we replotted peri-saccadic percept trajectories associated with rightward and leftward saccades (similar to Fig. 5A), but now while maintaining the original raw coordinates (Fig. 7A, B). We found that orthogonal mislocalization for off-axis flashes with nearer and farther components (i.e. #2, #4, #6, and #8) was much weaker for upper visual field flashes than for lower visual field ones, independently of whether a saccade was rightward or leftward (Fig. 7A, B). Lower visual field flashes showed consistent results with (Kaiser and Lappe 2004) in the sense that farther flashes had stronger orthogonal peri-saccadic mislocalization than nearer ones (Fig. 7A-F). Thus, for horizontal saccades, orthogonal perceptual mislocalization replicated (Kaiser and Lappe 2004) but only for lower visual field flash locations. These results are intriguing because they indicate that not only saccade vector direction matters for peri-saccadic perceptual mislocalization patterns (Figs. 1-6); retinotopic flash location also does matter (Fig. 7). Interestingly, closer inspection of the results of (Kaiser and Lappe 2004) with rightward saccades does indeed show that orthogonal mislocalization in their data was also weaker for upper visual field flashes than for lower visual field flashes, consistent with our results.

### Interaction between saccade direction vector and flash location for diagonal saccades

To further demonstrate that both saccade direction and flash location matter, we ran two of our subjects on a second experiment involving diagonal saccades (∼14.6 deg amplitude in the 45 deg rightward upward or rightward downward direction; Materials and Methods). Figure 8A, B shows the results of mislocalization in the same format as in Fig. 1. Two-dimensional peri-saccadic mislocalization was always aligned to the saccade vector (i.e. there was, in general, compression towards the saccade target along the saccade vector), and this is consistent with accounts of mislocalization relying on flash eccentricity relative to saccade target eccentricity (Hamker et al. 2008; Hamker et al. 2011; Kaiser and Lappe 2004; Richard et al. 2009; VanRullen 2004; Zirnsak et al. 2010). However, there were still differences in mislocalization as a function of saccade direction. For example, in Fig. 8C, we directly superimposed peri-saccadic percepts for diagonal upward and diagonal downward saccades using the same coordinate transformation (e.g. Fig. 2A). If neural maps responsible for peri-saccadic perceptual mislocalization were perfectly directionally symmetric (Hamker et al. 2008; Hamker et al. 2011; Kaiser and Lappe 2004; Richard et al. 2009; VanRullen 2004; Zirnsak et al. 2010), then the two contours shown in Fig. 8C should overlap. However, there were clear distortions, with diagonal upward saccades showing stronger mislocalization (also see Fig. 8D), especially for upper visual field flash locations (e.g. flashes #6 and #7; Supplementary Movie 1). Interestingly, for some flashes, diagonal saccades were associated with even stronger mislocalization than with upward saccades in the first experiment (Fig. 8E, F). Also, like in our first experiment, these results were evident even on the individual subject level (Fig. 9). Finally, when we plotted time courses of perceived flash positions for different flash locations in diagonal saccades, we found that the strongest mislocalizations (in the 0-50 ms peri-saccadic intervals) tended to occur when both the saccade direction and flash location had an upward component to them (Fig. 10). Thus, asymmetries in perceptual mislocalization for upward saccades compared to other cardinal directions also extended to cases with oblique saccades as well.

**Figure 10.**
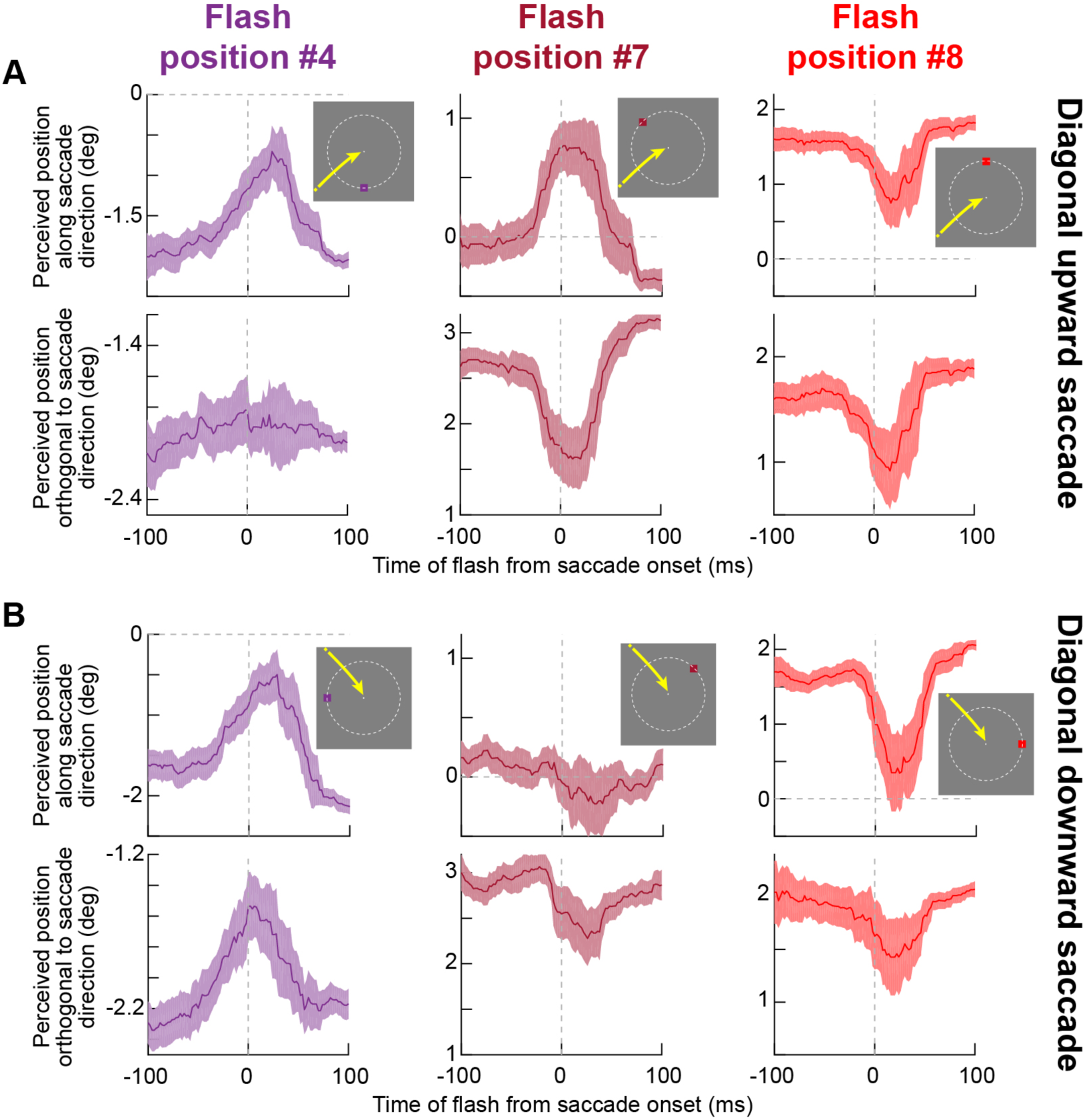
Time courses of perceptual mislocalization for diagonal upward and diagonal downward saccades revealed interactions between saccade direction and flash location in the phenomenon of peri-saccadic compression of space. (**A**) We plotted a time course of perceived flash position along (top row) or orthogonal to (bottom row) the axis of diagonal upward saccades for different flash locations (across columns). Error bars denote 95% confidence intervals. (**B**) Similar analysis for similar flash locations (relative to saccade direction and target location according to our standard convention), but now for diagonal downward saccades. Accounts of compression based on the retinotopic eccentricity of the flash relative to the saccade target location predict identical patterns of parallel (along the axis of the saccade) and orthogonal mislocalization between corresponding flash locations in **A** and **B**. However, there were obvious differences based on saccade direction. For example, flash #4 showed more orthogonal mislocalization for the diagonal downward saccade than the diagonal upward one. This pattern reversed between diagonal upward and downward saccades for flashes #7 and #8. Mislocalization was strongest when both the saccade and the flash locations had upper visual field locations (i.e. flashes #7 and #8). N=15-44 repetitions per time bin in the time courses.

### Implications for neural circuits involved in peri-saccadic mislocalization

In all, our results do not contradict the idea that foveal magnification plays a role in peri-saccadic perceptual mislocalization (Hamker et al. 2008; Hamker et al. 2011; Kaiser and Lappe 2004; Richard et al. 2009; VanRullen 2004; Zirnsak et al. 2010). However, these same results also suggest that the directional symmetry inherent in such accounts may be overly simplistic. To demonstrate this, we asked whether a simple neural circuit principle could make our results plausible. Our goal was not to exhaustively model and/or fit data, but rather to test the conceptual plausibility of the hypothesis that tissue asymmetries beyond foveal magnification can give rise to differential patterns of peri-saccadic mislocalization (also see **Discussion**). Our starting point was a common agreement among models (Hamker et al. 2008; Hamker et al. 2011; Kaiser and Lappe 2004; Richard et al. 2009; VanRullen 2004; Zirnsak et al. 2010) that neural maps with foveal magnification can account for mislocalization. While we are ambivalent to the details of individual models, we picked the simplest instantiation of them (VanRullen 2004), in which read-out of flash location is done after a simple translation of neural activity from the periphery (where the flash is presented) to the fovea (where the read-out of flash location takes place post-saccadically) (VanRullen 2004).

In the map of Fig. 11A (Materials and Methods), if there was a flash around an upward saccade target, then the flash would be around the magenta square in the figure (for example, one of the five shown flash locations). The saccade would translate this neural activity to the fovea (along the trajectory shown by the magenta arrow). The resulting percept would emerge by reading out the translated flash locations, which are shown by the magenta contour in the foveal representation of Fig. 11A (i.e. after the saccade). In Fig. 11A, we implemented such an idea on a map exhibiting not only foveal magnification (VanRullen 2004), but also an asymmetry between the representations of the upper and lower visual fields (Hafed and Chen 2016) (Materials and Methods). We also repeated the same exercise for horizontal (blue arrow) and downward (cyan arrow) saccades. Using this particular example map, based on physiological results from the SC (Hafed and Chen 2016), upward saccades (magenta arrow and contours in the foveal representation of Fig. 11A) would indeed result in foveal neural activity of flash locations (i.e. for read-out of the percept) that is an outlier when compared to all other cardinal saccade directions (other contours in the foveal region of the map in Fig. 11A). With a purely symmetric neural map exhibiting only foveal magnification (Fig. 11B) as in the scheme of (VanRullen 2004) and related models, none of the saccade directions would be outliers (Fig. 11B). Interestingly, a map like in Fig. 11A would also predict an asymmetry between upper and lower visual field flash locations even for purely horizontal saccades (e.g. Fig. 7) because of the asymmetry in the map itself.

**Figure 11.**
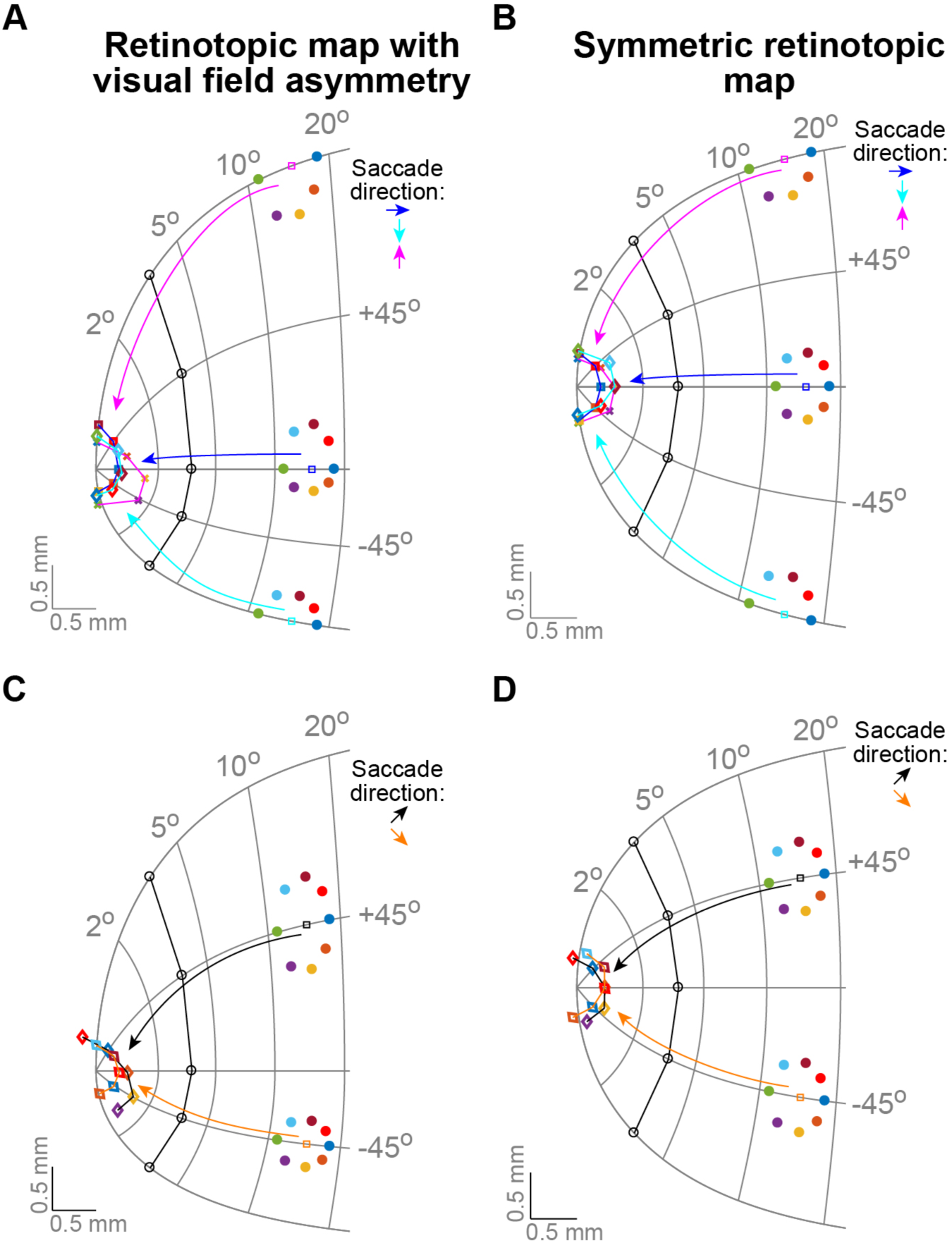
Asymmetries in peri-saccadic mislocalization can occur if sensory-motor visual maps have asymmetries in their upper and lower visual field representations. (**A**, **B**) It was suggested that patterns of two-dimensional mislocalization (e.g. Fig. 1A) can arise as a result of translation of neural activity, on a visual map containing foveal magnification, from the peripheral locus representing pre-saccadic target position to the foveal locus representing post-saccadic fixation. However, this account predicts similar mislocalizations regardless of saccade direction. We asked whether asymmetries in visual field representations can, theoretically, result in upward saccades having different peri-saccadic mislocalization. We simulated a map having both foveal and upper visual field magnification (**A**) or a map having only foveal magnification but symmetric upper/lower visual fields (**B**). The equations for the maps are cited in Materials and Methods. Also, the maps are oriented such that eccentricity is organized along the horizontal axis of neural tissue (curved vertical lines indicate iso-eccentricity loci) and direction from horizontal is organized along the vertical axis (curved horizontal lines indicate iso-direction loci; e.g. +45° and −45°). Horizontal or vertical saccades (squares), as well as their associated flash locations (colored circles), translate neural activity associated with the flashes towards the foveal origin of the map along the saccade trajectory (colored arrows). We used a simple translation along the axis of each saccade direction in the neural tissue. Because of foveal magnification, the resulting neural activity translated from the periphery (colored lines in the foveal region) is mislocalized relative to the veridical flash locations (black lines around the foveal region). Notice how a symmetric map (**B**) predicts similar mislocalization patterns for all saccade directions, whereas an asymmetric map (**A**) predicts that upward saccades would be outliers. (**C**,**D**) Similarly, a symmetric map (**D**) predicts similar mislocalization patterns for diagonal upward versus downward saccades, but an asymmetric map (**C**) is more consistent with our observations of a difference in mislocalization between these otherwise-identical saccades.

Similarly, when simulating the same translation mechanism on the asymmetric map for diagonal saccades (Fig. 11C), then percepts associated with diagonal upward versus diagonal downward saccades would be distorted with respect to each other (Fig. 11C), just like in our data (Figs. 8-10). This would again not be the case with a purely symmetric map (Fig. 11D). Thus, an asymmetry beyond foveal magnification in neural maps is a plausible mechanism for our observation that peri-saccadic perceptual mislocalization is different for upward saccades.

## Discussion

We observed a difference in peri-saccadic perceptual mislocalization for upward saccades when compared to other cardinal saccade directions. We also noticed that even for purely horizontal saccades, upper versus lower visual field retinotopic flash locations experience different patterns of peri-saccadic perceptual mislocalization. We think that these results are particularly interesting because they touch on a broader question of how different sensory, sensory-motor, and motor maps in the brain interact around the time of saccades in order to ensure trans-saccadic perceptual stability. Specifically, if peri-saccadic perceptual mislocalization really does depend on neural tissue distortions (Hamker et al. 2008; Hamker et al. 2011; Kaiser and Lappe 2004; Richard et al. 2009; VanRullen 2004; Zirnsak et al. 2010), then a significant question is: what happens if some neural maps involved in the phenomenon have additional asymmetries (beyond foveal magnification) that are not present in other maps functionally connected to them at the time of saccades? For example, if the SC has upper visual field magnification (Hafed and Chen 2016) whereas the primary visual cortex (V1) or area V4 not, then how might these different neural maps interact with each other during peri-saccadic intervals if the maps were critical for creating the perceptual phenomenon? Similarly, if some areas have upper/lower visual field asymmetries and others not, then perhaps observations like ours can pinpoint potential neural loci for peri-saccadic perceptual mislocalization.

Such loci are not necessarily known, and there is an urgent need to explore them in order to resolve some of the ongoing debates in the literature about perceptual stability. For example, the frontal eye fields (FEF) and V4 exhibit peri-saccadic response field (RF) changes that have been implicated in peri-saccadic perceptual mislocalization (Hamker et al. 2008; Hamker et al. 2011; Tolias et al. 2001; Zirnsak et al. 2010; Zirnsak et al. 2014), although with some debate on the exact details of the mechanisms (Hartmann et al. 2017; Neupane et al. 2016a; b). Moreover, even if there were no such debates, it is not known whether FEF or area V4 do exhibit upper/lower visual field asymmetries like the ones recently described in the SC (Hafed and Chen 2016) or not. It is thus not yet known how our experimental results above can emerge neurophysiologically. It would be highly interesting to use our results to motivate neurophysiological investigations that can pinpoint neural loci for peri-saccadic perceptual mislocalization phenomena, and to specifically ask whether the SC is indeed relevant for these phenomena or not. These investigations must also pair neural recordings with behavior (in the same animals) to make the most sense. For example, in the other peri-saccadic phenomenon of saccadic suppression alluded to in **Introduction**, only when linking neurons to behavior in the same animals have certain perceptual properties of saccadic suppression found a convincing neural locus (Chen and Hafed 2017).

Having said the above, it should be emphasized that our results and model so far do not unequivocally implicate the SC in peri-saccadic perceptual mislocalization, although we do think that the SC might be involved given the details of our results. In our model and experiments, we simply used the SC (Hafed and Chen 2016) as motivation for the idea that asymmetries in neural maps for upper and lower visual field representations can be relevant for peri-saccadic perceptual mislocalization, and this was sufficient to uncover the observations that we have documented above. Neurophysiological experiments would be critically needed to investigate whether the SC is indeed relevant and how.

Theoretically speaking, maybe even the asymmetry in relevant neural maps can be in the opposite direction from that in the SC and our model but still result in upward saccades being outliers with appropriate read-out mechanisms. However, we think that results like those in Fig. 7 would suggest that the perceptual phenomenon likely does depend on maps with compressed lower rather than upper visual field representations. That is, if distortions in percepts for farther flash locations are due to compressed neural tissue at large eccentricities (Hamker et al. 2008; Hamker et al. 2011; Kaiser and Lappe 2004; Richard et al. 2009; VanRullen 2004; Zirnsak et al. 2010), which increases uncertainty about true flash location, then the presence of orthogonal perceptual distortions for lower rather than upper visual field flash locations in Fig. 7 would argue, by the same logic, that the perceptual phenomenon depends on neural maps with compressed neural tissue for lower visual field locations, like in the SC (Hafed and Chen 2016).

It must also be emphasized that models other than the translation model, which we have highlighted as an example in Fig. 11, can also account for our results if asymmetries between upper and lower visual fields are introduced. Specifically, in such models (Hamker et al. 2008; Hamker et al. 2011; Kaiser and Lappe 2004; Richard et al. 2009; Zirnsak et al. 2010), mislocalization occurs because of an interaction between an oculomotor command in oculomotor maps and a visual response in visual maps. According to these models, specific patterns of mislocalization (e.g. Fig. 1A) arise, at least in part, because both the oculomotor and visual maps that are interacting exhibit foveal magnification. If these maps were to also exhibit other tissue asymmetries beyond foveal magnification, such as upper and lower visual field asymmetries, then interactions between them can still potentially account for our results. It would be interesting to understand how different maps exhibiting different amounts or types of magnification would need to be connected together in order to account for the asymmetries that we have observed in our current experiments.

Another intriguing aspect of our results is again related to Fig. 7, for which we found results very different from those in (Kaiser and Lappe 2004) for horizontal saccades (at least for some flash locations). Specifically, we found that upper visual field flashes did not experience orthogonal mislocalization whereas lower visual field flashes did. While we do think that close inspection of their data reveals an asymmetry between upper and lower visual field flash locations consistent with our observations, it may be asked why the difference in Fig. 7 that we saw between upper and lower visual fields was so dramatic compared to their results. One possibility is that our saccades were slightly smaller than those used in (Kaiser and Lappe 2004). With larger saccades, flashes and saccade targets are at more eccentric neural loci than with smaller saccades, which means neural tissue representing saccade and flash eccentricities is even more compressed when compared to the case with smaller saccades. This naturally magnifies the mislocalization according to existing models (Hamker et al. 2008; Hamker et al. 2011; Kaiser and Lappe 2004; Richard et al. 2009; VanRullen 2004; Zirnsak et al. 2010). Thus, with our smaller saccades, we may have sampled different neural loci than those by (Kaiser and Lappe 2004). It would be interesting to extend our and their results to different saccade amplitudes. Indeed, peri-saccadic perceptual mislocalization still takes place for small saccades (Lavergne et al. 2010) and microsaccades (Hafed 2013).

Finally, we think that it would be interesting to explore reasons behind certain other observations in our data, like the early bias in percepts for some flash locations in Fig. 5 well before any peri-saccadic transient changes in perception started to occur. These biases, and our results overall, relate to asymmetries in several aspects of visual-motor behavior (Greenwood et al. 2017; Hafed and Chen 2016; He et al. 1996; Rubin et al. 1996), and understanding the commonalities and differences between these different phenomena would be important for pinpointing at which stage of visual processing peri-saccadic perceptual mislocalization phenomena start to take place.

## Acknowledgments

We were funded by the Werner Reichardt Centre for Integrative Neuroscience (CIN) at the Eberhard Karls University of Tübingen. The CIN is an Excellence Cluster funded by the Deutsche Forschungsgemeinschaft (DFG) within the framework of the Excellence Initiative (EXC 307). We were also funded by the Hertie Institute for Clinical Brain Research at the Eberhard Karls University of Tübingen.

## Supplementary Movie

**Supplementary Movie 1 Demonstration of the dependence of peri-saccadic perceptual mislocalization on saccade direction for diagonal upward versus diagonal downward saccades.** The movie shows the instantaneous perceived position of the different flashes around the saccade target, using the same running window averages employed for the time course analyses of Figs. 5, 7, 8, 10. Percepts associated with diagonal upward saccades are displayed in black, and percepts associated with diagonal downward saccades are displayed in orange. Both saccade directions were rotated as in all of our analyses such that horizontal displacements in the movie reflect mislocalizations along the direction of the saccade vector, and vertical displacements reflect mislocalizations orthogonal to the direction of the saccade vector. In other words, the data were rotated as if both conditions were associated with a rightward saccade vector in the movie. Flash locations around the saccade target are color-coded using the same conventions described in Fig. 1. Note how the percept is not simply a function of eccentricity of the flash relative to saccade target eccentricity, in which case mislocalizations should be similar for all flash locations between the two saccade directions. Instead, there were asymmetries in percepts associated with upward-directed and downward-directed saccades.

